# N-Acetyl Cysteine Abrogates Silver-Induced Reactive Oxygen Species in Human Cells Without Altering Silver-based Antimicrobial Activity

**DOI:** 10.1101/792150

**Authors:** Kush N. Shah, Parth N. Shah, Andrew R. Mullen, Qingquan Chen, Ralph J. DeBerardinis, Carolyn L. Cannon

## Abstract

Silver-based antimicrobials are widely used topically to treat infections associated with multi-drug resistant (MDR) pathogens. Expanding this topical use to aerosols to treat lung infections requires understanding and preventing silver toxicity in the respiratory tract. A key mechanism resulting in silver-induced toxicity is the production of reactive oxygen species (ROS). In this study, we have verified ROS generation in silver-treated bronchial epithelial (16HBE) cells prompting evaluation of three antioxidants, N-acetyl cysteine (NAC), ascorbic acid, and melatonin, to identify potential prophylactic agents. Among them, NAC was the only candidate that abrogated the ROS generation in response to silver exposure resulting in the rescue of these cells from silver-associated toxicity. Further, this protective effect directly translated to restoration of metabolic activity, as demonstrated by the normal levels of citric acid cycle metabolites in NAC-pretreated silver-exposed cells. As a result of the normalized citric acid cycle, cells pre-incubated with NAC demonstrated significantly higher levels of adenosine triphosphate (ATP) levels compared with NAC-free controls. Moreover, we found that this prodigious capacity of NAC to rescue silver-exposed cells was due not only to its antioxidant activity, but also to its ability to directly bind silver. Despite binding to silver, NAC did not alter the antimicrobial activity of silver.

**Importance:** Although silver is a potent, broad-spectrum antibiotic, silver-induced toxicity, primarily due to generation of ROS, remains a concern limiting its use beyond treatment of wounds. NAC has been widely used as an antioxidant to rescue eukaryotic cells from metal-associated toxicity. Thus, we have evaluated the capacity of NAC to abrogate silver toxicity in a human bronchial epithelial cell line (16HBE) as a step towards expanding the use of silver-based antimicrobials to treat lung infections. We found that NAC pre-incubation resurrects a healthy metabolic state in bronchial epithelial cells exposed to silver ions via a combination of its antioxidant and metal-binding properties. Finally, this ability of NAC to rescue silver-exposed eukaryotic cells does not alter the antimicrobial activity of silver. Thus, a silver-NAC combination holds tremendous potential as a future, non-toxic antimicrobial agent.

## Introduction

Silver is a mainstay therapeutic strategy for prophylaxis, as well as eradication, of established infections in wound and burn patients (1). This wide-spread use of silver stems from its broad-spectrum antimicrobial activity and multiple mechanisms of action including disruption of bacterial cell walls, and DNA condensation (2, 3). These multiple mechanisms impart potent biocidal activity against several bacterial pathogens including multi-drug resistant (MDR) *Pseudomonas aeruginosa*, *Staphylococcus aureus, Escherichia coli*, as well as fungus, mold, and yeast (2, 4, 5). The ability of silver to target multiple pathways also lowers the propensity of resistance acquisition by microbes, which is commonly observed among antibiotics with single targets (2–5). Only a few cases of silver resistance have been reported (6). Thus, silver has been incorporated into, or used as a coating in, over 400 medical and consumer products including wound dressings, catheters and endotracheal tubes, bone cement, socks, and disinfectants (7). In addition to its antimicrobial activity, silver has also garnered attention as a potential anticancer agent (8). Despite this tremendous potential, stability and toxicity are two major limitations that hamper the use of silver as a therapeutic on a larger scale.

The oligodynamic effects of silver are limited to its ionic form (+1 oxidation state; Ag^+^), which has a high affinity for chloride ions, as well as thiol functionalized substrates and proteins (6, 9). Interaction with these functional groups often results in deactivation of the silver ion and loss of biological activity (6). Cannon and Youngs have developed a library of silver-based antimicrobials, silver carbene complexes (SCCs), with enhanced stability over conventional silver salts (10–14). These molecules have demonstrated superior antimicrobial activity against clinically relevant MDR pathogens including *Pseudomonas aeruginosa*, *Burkholderia cepacia* complex species, *Staphylococcus aureus*, *Klebsiella pneumoniae*, and *Acinetobacter baumannii*, both *in vitro* and *in vivo* (10–16). These compounds also demonstrate potent antimicrobial activity against biodefense pathogens *Bacillus anthracis* and *Yersinia pestis* (15). Further, polymeric nanoparticles loaded with these SCCs demonstrate superior *in vivo* antimicrobial activity over parent molecules (15, 17). These devices offer sustained release of the therapeutic at the infection site and protect the silver ions from deactivation. As a result, in an acute pneumonia model, mice treated with SCC-loaded nanoparticles exhibit increased survival and lower bacterial burden with fewer and lower doses compared with treatment with unencapsulated SCCs (17). Thus, development of novel molecules and delivery devices have addressed the stability concerns and significantly improved the efficacy of silver, opening up new avenues for the use of silver beyond topical therapy.

Toxicity of silver has always been a controversial topic. Several publications report silver to be non-toxic, with argyria, a rare and irreversible pigmentation of the skin caused by silver deposition, as the only reported side-effect (6, 18). On the other hand, several reports have demonstrated toxic side effects of silver in eukaryotic cells; claims that are underscored by the anticancer activity of silver. While silver toxicity and chemotherapeutic activity have been reported, little is known about the molecular mechanisms that contribute to silver toxicity. Recently, several reports have focused on identifying the mechanisms that contribute to toxicity towards eukaryotic cells, and are also responsible for the anticancer activity of silver nanoparticles (5, 18–25). These reports largely focus on the effect of size and surface coatings on toxicity of metallic silver nanoparticles (19–27). In general, pure silver nanoparticles demonstrate lower toxicity to eukaryotic cells compared with ionic silver at comparable concentrations (27), likely resulting from the gradual release of ionic silver from the nanoparticles upon surface oxidation or dissolution. Kittler *et al.* (26) have established a direct correlation between dissolution of nanoparticles and subsequent release of silver ions to toxicity towards eukaryotic cells. While the individual toxicity caused by nanoparticles and silver ions has yet to be discerned, generation of reactive oxygen species (ROS) has been implicated as a key underlying mechanism of toxicity in both instances (28, 29). ROS and the complementary cellular antioxidant defense system are part of a complex cellular milieu that plays critical roles in several biochemical processes (30). Silver disrupts the mitochondrial respiratory chain resulting in overproduction of ROS, leading to oxidative stress, ultimately causing lipid peroxidation and protein denaturation, interruption of ATP production, DNA damage, and induction of apoptosis (31, 32). Thus, ROS overproduction is one of the primary mechanisms responsible for inhibition of cell proliferation and induction of cell death in cells exposed to silver. N-acetyl cysteine (NAC) has been employed as an antioxidant to abrogate ROS generation and alleviate toxicity of silver towards eukaryotic cells (29, 33–35). However, the effects of anti-oxidants such as NAC on the overall cellular health and cell metabolism are not well known.

We aim to develop non-toxic therapeutic strategies for eradication of multi-drug resistant bacterial pathogens, particularly, pathogens responsible for lung infections. We have extensively demonstrated the antimicrobial activity of silver against several pathogens that result in lung infections, and here we evaluate the impact of silver-based compounds on host cellular metabolism. Because we are interested in developing silver-based antimicrobials to treat lung infections, we have evaluated the toxicity of silver in a human bronchial epithelial cell line (16HBE). These studies are also relevant to environmental inhalation exposures to silver particulates. We first confirmed that silver induces ROS in these 16HBE cells. Next, we have determined the effect of three antioxidants, ascorbic acid (vitamin C), melatonin, and NAC, on cell viability, and identified NAC as the only antioxidant that results in reduction of silver toxicity. Further, we demonstrate the ability of NAC to reduce ROS generation in these cells without affecting glutathione concentrations. Finally, exposure to silver disrupts cellular metabolism; pre-incubation with NAC rescues the cells allowing maintenance of ATP production. Additionally, the ability of NAC to rescue cells from silver toxicity is not limited to 16HBEs; we have demonstrated similar effects in human dermal fibroblasts. Thus, NAC pre-incubation suppresses ROS generation and maintains metabolic activity of the cell by sequestering silver ions to abrogate silver toxicity.

## Results

### Silver induction of reactive oxygen species (ROS) and superoxide

Several publications have demonstrated ROS generation by eukaryotic cells after exposure to silver. Thus, we sought to verify the observation that silver acetate induces reactive oxygen species and superoxide ions in a human bronchial cell line, 16HBE. Our results demonstrated a significantly higher amount of ROS and superoxide ions within cells that are incubated with silver acetate, at 8 and 24 hours, compared with cells that are not exposed to silver (Figure 1).

**Figure 1.**
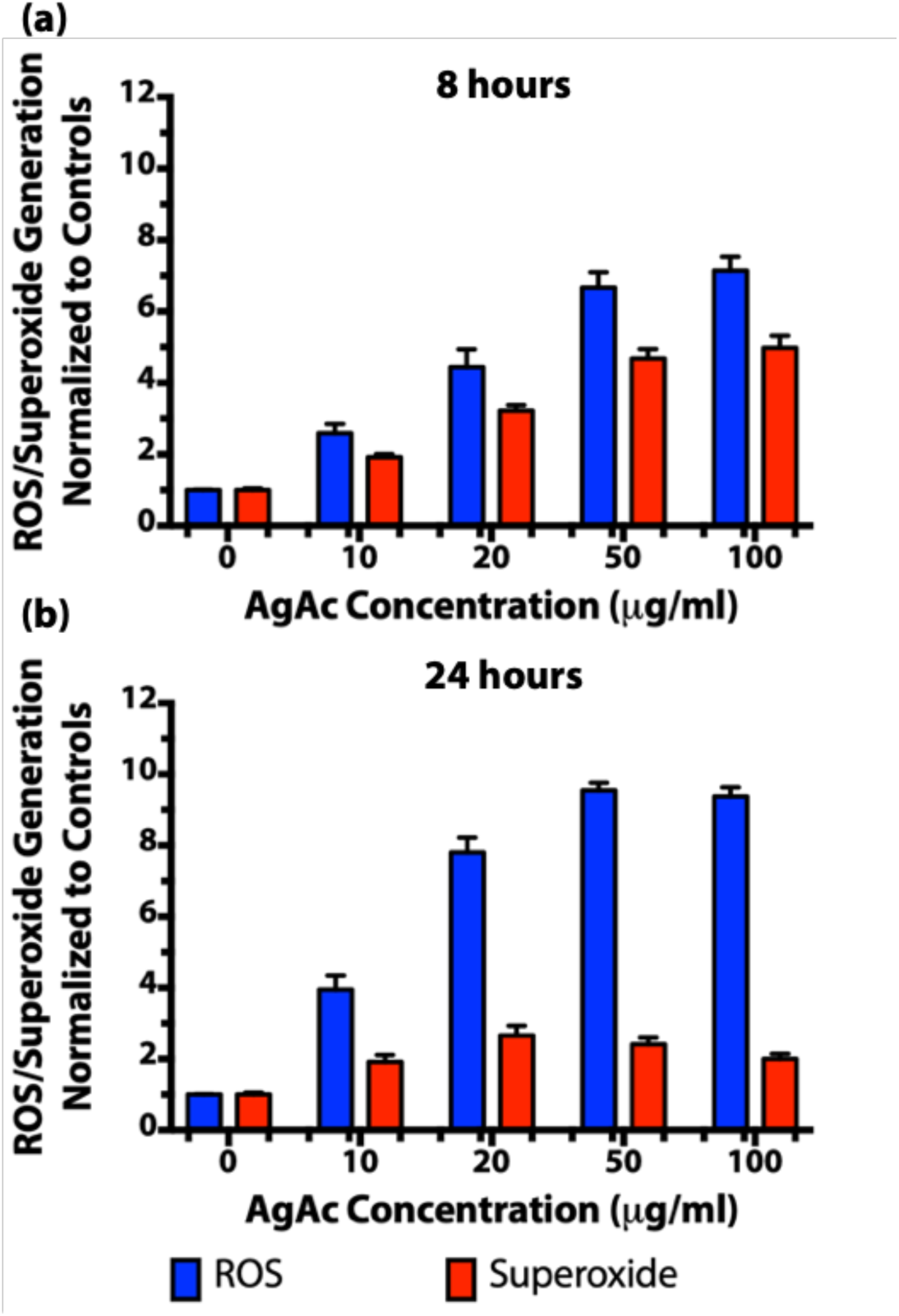
Reactive oxygen species and superoxide levels in human bronchial epithelial (16HBE) cells upon exposure to silver acetate for (a) 8 and (b) 24h.

### Activity of antioxidants

Comparing the antioxidant activity of NAC, ascorbic acid, and melatonin against the antioxidant standard, Trolox, a water-soluble vitamin E analog, in a cell-free assay, demonstrated ascorbic acid to have the highest antioxidant capacity, followed by NAC (Figure 2). Ascorbic acid demonstrated significantly higher antioxidant activity compared with melatonin and NAC (*p* ≤ 0.0001); NAC demonstrated significantly higher antioxidant activity compared with melatonin (*p* ≤ 0.001). This superior antioxidant capacity of ascorbic acid was also evident in cells incubated with ascorbic acid. Cells incubated with ascorbic acid demonstrated significantly higher antioxidant activity over cells only, and NAC incubated cells (*p* ≤ 0.01), but not melatonin incubated cells. Surprisingly, NAC incubation did not result in enhanced anti-oxidant activity, as no significant difference was observed between NAC incubated cells and melatonin incubated cells, or cells incubated with media only.

**Figure 2.**
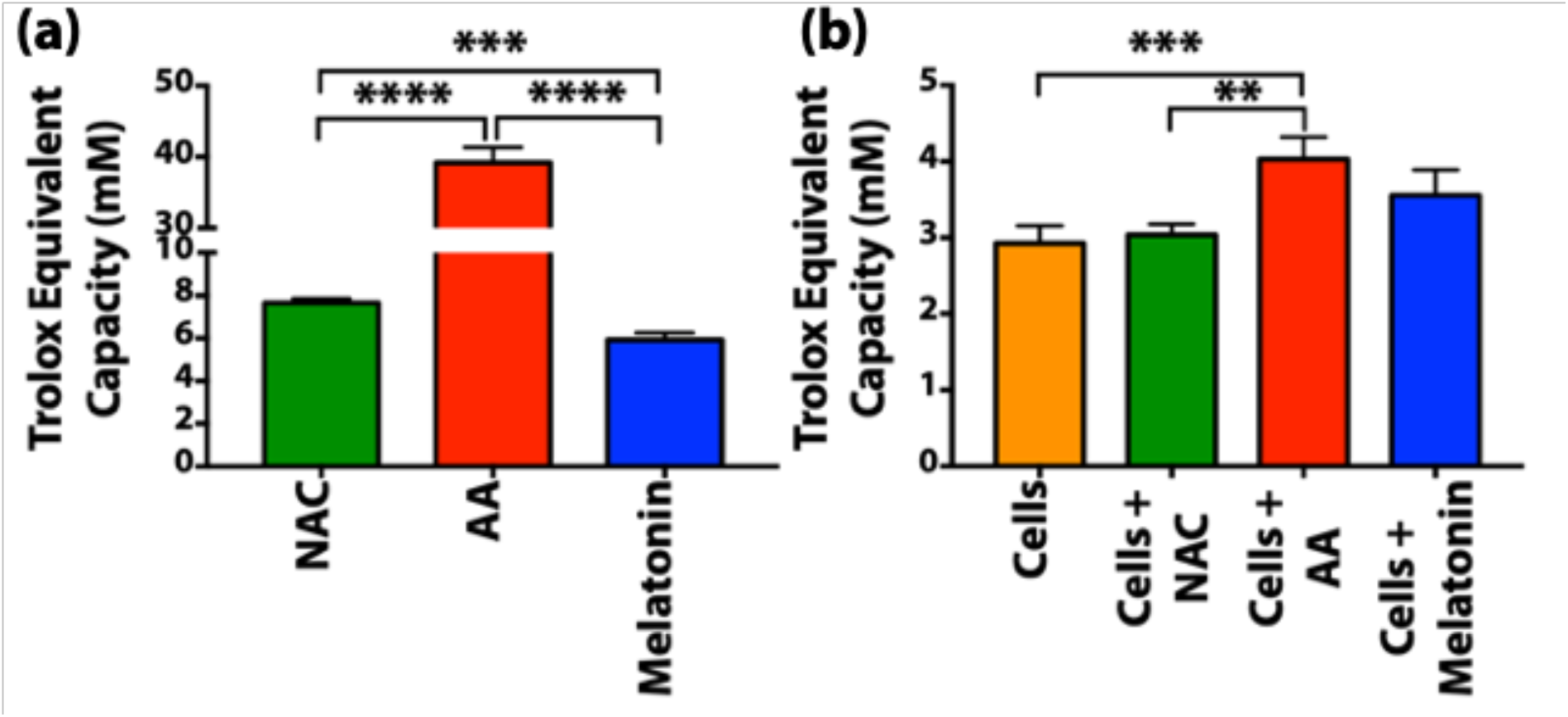
Antioxidant activity of (a) N-acetyl cysteine (NAC), ascorbic acid (AA), and melatonin at 10 mM concentration, and (b) cell lysates from cells incubated with no antioxidant, 10 mM N-acetyl cysteine, 10 mM ascorbic acid or 10 mM Melatonin for two hours. **: *p* ≤ 0.01, ***: *p* ≤ 0.001, and ****: *p* ≤ 0.0001.

### Abrogation of silver acetate toxicity through pre-incubation with antioxidants

The effect of antioxidant pre-incubation on silver acetate toxicity towards 16HBE cells is shown in Figure 3 (and Figure S1 in supplementary information). Cells pre-incubated with NAC demonstrated significantly higher survival upon exposure to silver acetate at 8 and 24 hours. Cells pre-incubated with 10 mM NAC and up to 100 μg/mL silver acetate demonstrated significantly higher survival over cells that were not pre-incubated with NAC (*p* ≤ 0.05). Similar results were observed with cells pre-incubated with 5.0 and 7.5 mM NAC followed by incubation with silver acetate at 50 and 75 μg/mL, respectively. Moreover, a dose response was observed; NAC pre-incubation at concentrations 1.0 mM or higher result in lower silver acetate toxicity. The lethal dose at median cell viability (LD_50_) values for cells exposed to silver acetate after pre-incubation with 1, 2.5, 5, 7.5, and 10 mM NAC were 16, 27, 36, 63, and 60 μg/mL respectively, in contrast to a LD_50_ of 7.8 μg/mL silver acetate when cells were not pre-incubated with NAC.

**Figure 3.**
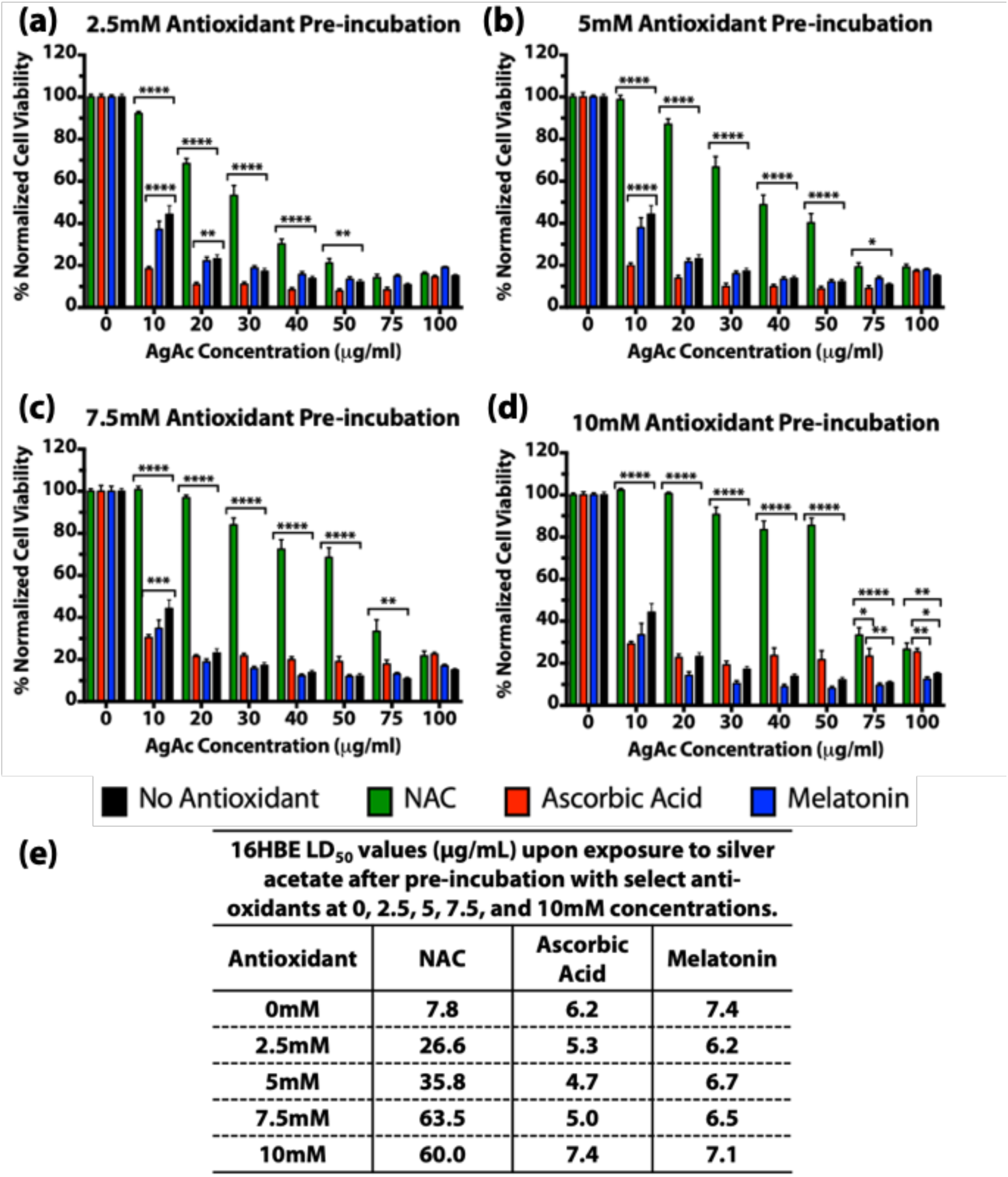
Viability of human bronchial epithelial (16HBE) cells upon pre-incubation with 0, 2.5, 5, 7.5, and 10 mM concentrations of N-acetyl cysteine (NAC), ascorbic acid, or melatonin for 2 h followed by a 24 h incubation with 0, 10, 20, 30, 40, 50, 75, or 100 μg/mL concentration of silver

Pre-incubation with neither ascorbic acid nor melatonin resulted in increased cellular survival comparable to NAC. Pre-incubation with 7.5 and 10 mM ascorbic acid followed by up to 100 μg/mL silver acetate exposure resulted in significantly higher cell survival (*p* ≤ 0.05). Despite the higher survival, however, the LD_50_ values upon pre-incubation with ascorbic acid did not appreciably change. Moreover, cells exposed to 50 μg/mL silver acetate after 10 mM NAC pre-incubation exhibited 86% survival compared with 22% survival upon pre-incubation with 10 mM ascorbic acid (*p* ≤ 0.0001). Finally, melatonin pre-incubation did not alter the toxicity of silver acetate as demonstrated by the cell survival and LD_50_ values. Thus, of the three anti-oxidants evaluated in this study, only NAC rescued the cells from silver acetate toxicity. Additionally, the CyQuant cell viability assay demonstrated similar results with 0 mM and 10 mM NAC pre-incubation (Figure S2, supplementary information). The CyQuant cell viability assay also demonstrated the onset of silver acetate toxicity after only one hour of incubation. Finally, NAC pre-incubation also abrogated the toxicity of silver acetate against human dermal fibroblasts (Figure S3, supplementary information). Thus, NAC was chosen as the molecule of interest for further investigation.

### Silver induction of reactive oxygen species (ROS) and superoxide

Figure 4 illustrates the effect of NAC on silver acetate induced reactive oxygen species and superoxide ions. Pre-incubation with NAC suppressed the levels of ROS and superoxide seen after incubation with silver acetate for 8 and 24 hours. Specifically, cells pre-incubated with 10 mM NAC, upon exposure to silver acetate concentrations higher than 20 μg/mL, showed significantly lower ROS levels at 8 and 24 hours (*p* ≤ 0.001). Similarly, when cells were pre-incubated with 10 mM NAC, superoxide levels were significantly lower at 8 and 24 hours after incubation with 10, 20, and 50 μg/mL silver acetate (*p* ≤ 0.001). Cells incubated with 100 μg/mL silver acetate showed significantly lower superoxide levels at 8 hours when pre-incubated with NAC (*p* ≤ 0.01), but not at 24 hours. Surprisingly, NAC pre-incubation initially induced ROS, which subsided after 8 hours, but had no effect on superoxide levels.

**Figure 4.**
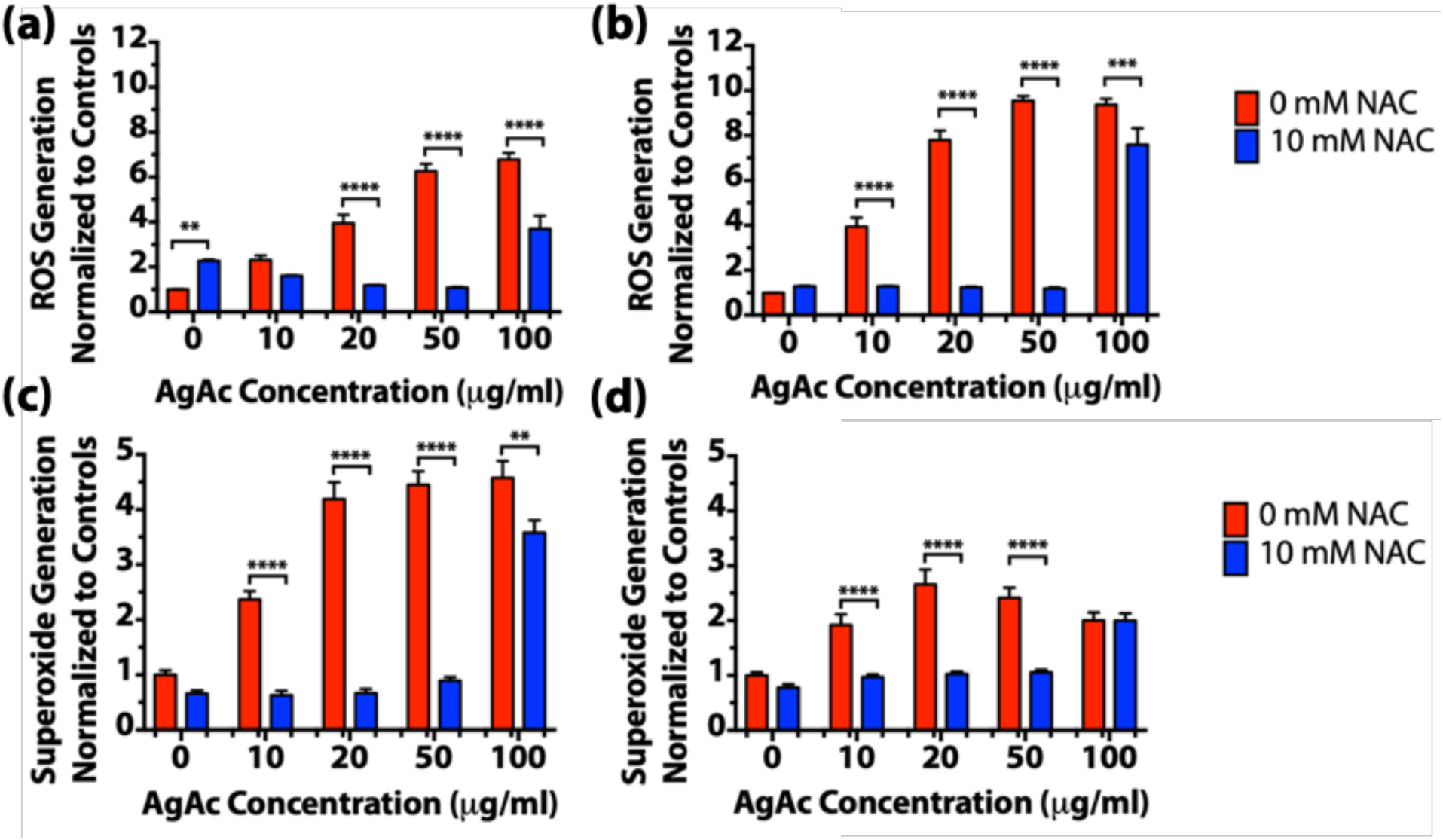
Reactive oxygen species (a, b) and superoxide (c, d) levels in human bronchial epithelial (16HBE) cells upon pre-incubation with 0 or 10 mM NAC followed by a (a, c) 8 h or (b, d) 24 h exposure to silver acetate. **: *p* ≤ 0.01 and ****: *p* ≤ 0.0001.

### Glutathione concentrations after treatment with NAC

Because NAC is a known precursor of glutathione, the effect of NAC pre-incubation on both oxidized and reduced glutathione concentrations was determined. A dose response and significant reduction in the reduced glutathione concentration was observed with increasing concentrations of silver, with or without NAC pre-treatment (Figure 5). However, this increase in GSH concentration was not accompanied by an increase in GSSG levels, suggesting the absence of correlation between ROS generation and oxidation of glutathione, after silver incubation. Further, NAC pre-treatment did not result in any changes in total, oxidized, or reduced glutathione concentrations over cells not pre-treated with NAC, with or without an insult from silver.

**Figure 5.**
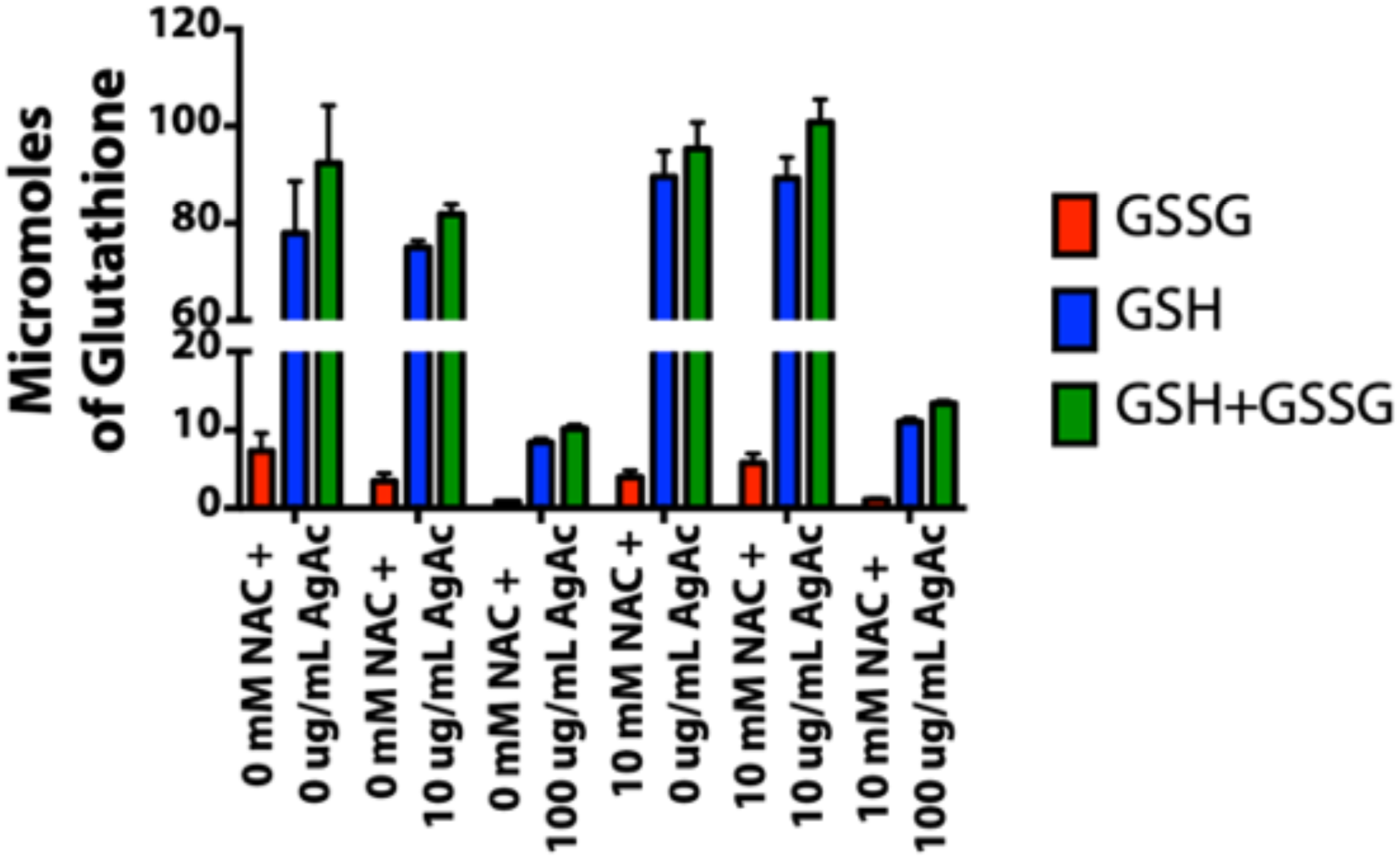
Levels of reduced (GSH), oxidized (GSSG), and total (GSH+GSSG) glutathione measured in human bronchial epithelial (16HBE) cells upon pre-incubation with 0 or 10 mM NAC followed by exposure to 0, 10, or 100 μg/mL silver acetate (AgAc) for 1 h.

### Analysis of total metabolite pool size and metabolite labeling patterns using gas chromatography-mass spectroscopy

Disruption of the mitochondrial electron transport chain has been linked to the ROS overproduction and cell death upon exposure to silver ions. To further explore the metabolic effects of silver-induced ROS production, we evaluated glucose consumption and its metabolism through the glycolysis pathway. Glucose consumption and lactate production, the end product of glycolysis, were determined using a bioProfile BASIC analyzer (Figure 6). No significant difference was observed in glucose consumption and lactate production between cells that were pre-treated with 0 and 10 mM NAC; thus, incubation with NAC did not significantly alter glucose consumption and lactate production. As expected, treatment with increasing concentrations of silver acetate resulted in reduced glucose consumption and lactate production. Next, we evaluated the effect of NAC incubation on the oxidation of glucose-derived carbon in the TCA cycle. Levels of metabolites involved in the TCA cycle were evaluated using GC-MS and normalized to control cells (not exposed to silver or NAC; Figure 7). Cells exposed to silver alone demonstrated no change in lactate levels, but significantly lower levels of TCA cycle intermediates including citrate, fumarate, and malate, as well as glutamate and aspartate, surrogates for α-ketoglutarate and oxaloacetate, respectively, while silver-exposed cells pretreated with NAC showed significantly less attenuation of TCA intermediates. In particular, citrate, glutamate, fumarate, and malate levels were significantly higher for NAC pre-incubated cells after exposure to 30, 40, 50, and 75 μg/mL silver acetate (*p* ≤ 0.05). Aspartate levels were significantly higher for NAC pre-incubated cells upon exposure to 50 and 75 μg/mL silver acetate (*p* ≤ 0.05). Finally, lactate levels were significantly higher for cells pre-incubated with NAC and exposed to 50 μg/mL silver acetate only. In addition, pre-incubation with NAC does not appreciably alter the labeling patterns of key metabolites (Figure S4, supplementary information). Thus, these results demonstrated that exposure to silver acetate resulted in mitochondrial stress that can be ameliorated by pre-incubation with NAC.

**Figure 6.**
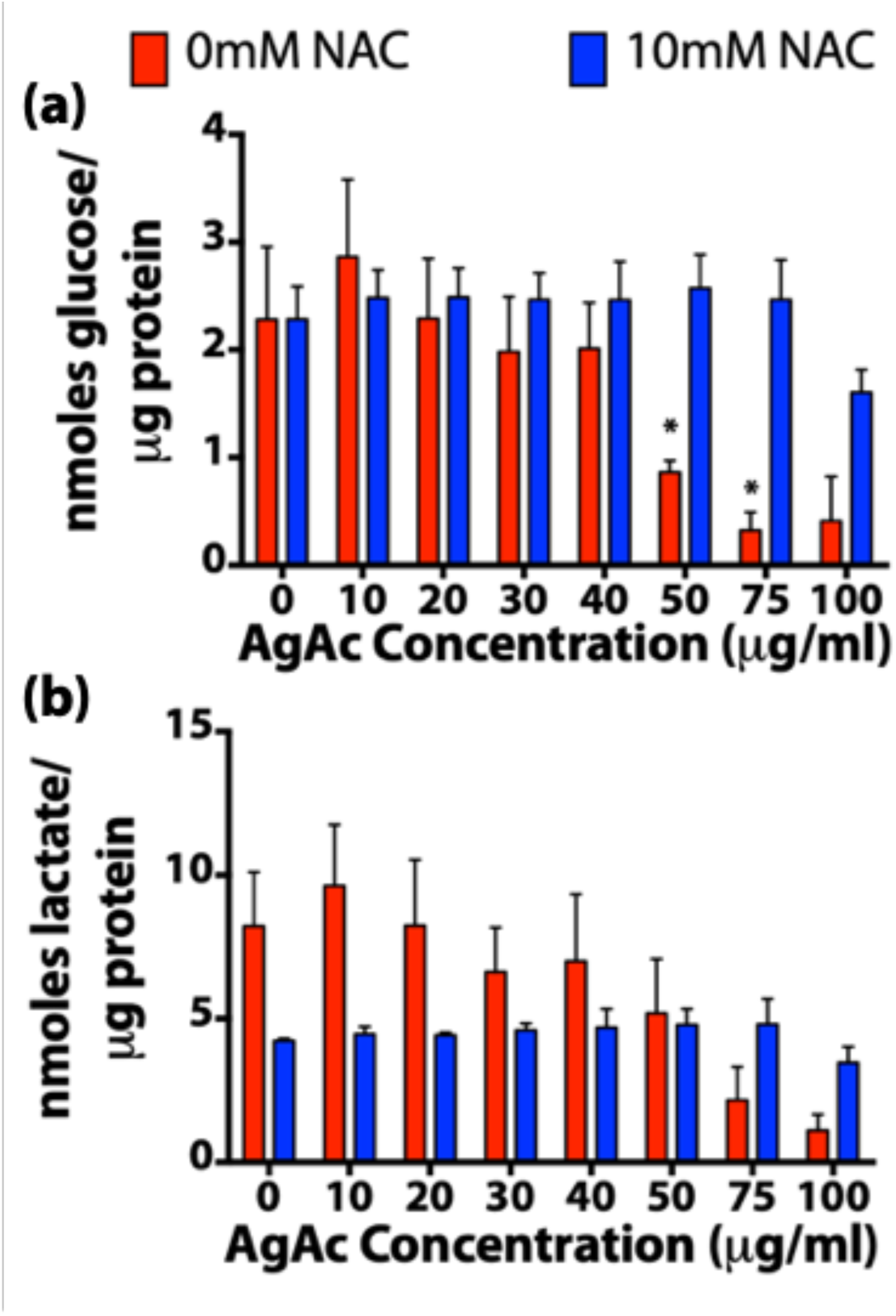
Levels of (a) glucose consumption and (b) lactate production in human bronchial epithelial cells (16HBE) upon pre-incubation with 0 or 10 mM N-acetyl cysteine (NAC) followed by an 8 h exposure to silver acetate, determined using a BioProfile BASIC analyzer. *: *p* ≤ 0.05.

**Figure 7.**
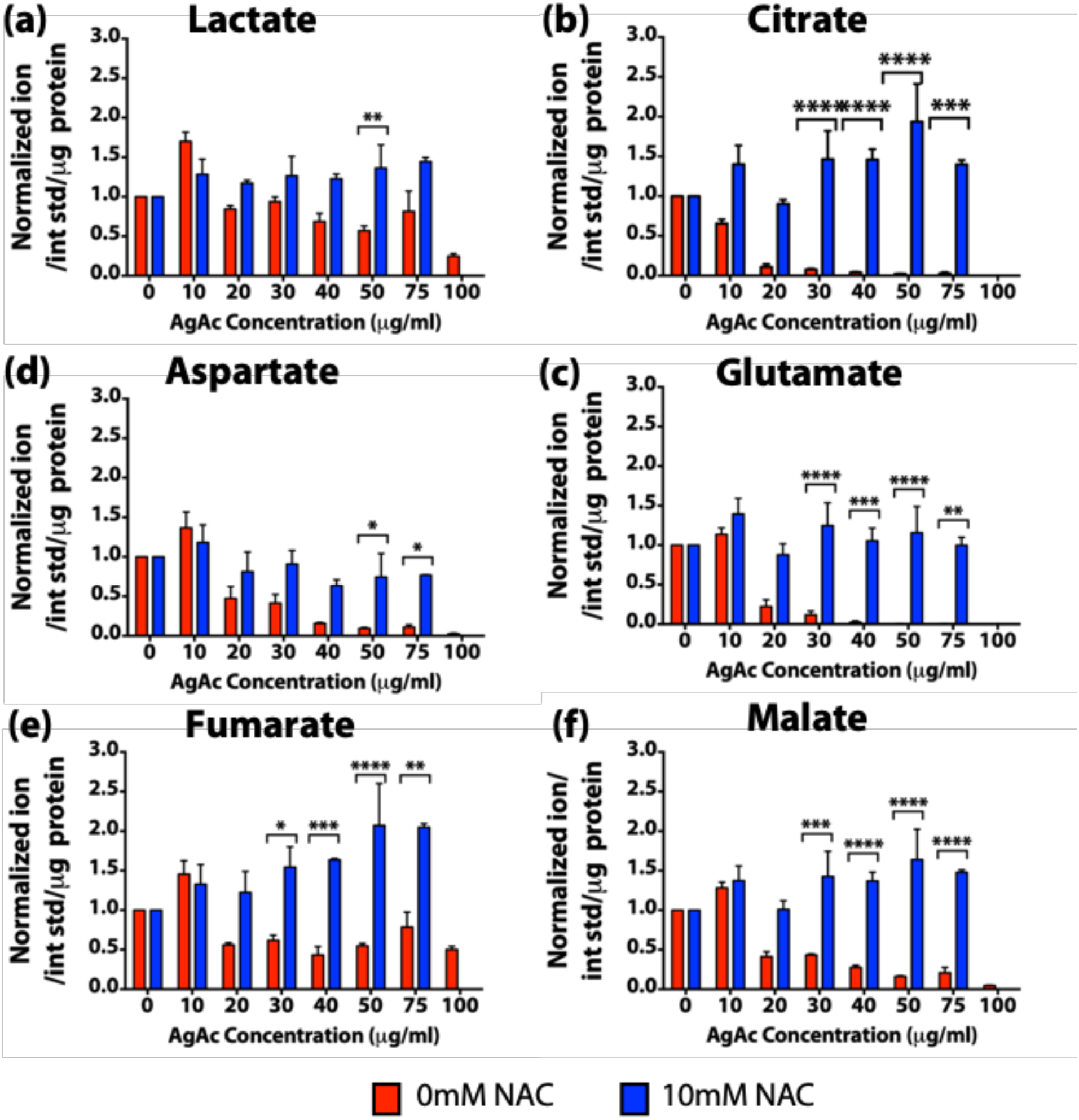
Effect of N-acetyl cysteine (NAC) pre-incubation (2h) on the TCA cycle metabolite pool after exposure to silver (8 h) in human bronchial epithelial (16HBE) cells; Levels of (a) lactate, (b) citrate, (c) glutamate, (d) aspartate, (e) fumarate, and (f) malate in the cell lysate determined using tandem gas chromatography – mass chromatography. *: *p* ≤ 0.05, **: *p* ≤ 0.01, ***: *p* ≤ 0.001, and ****: *p* ≤ 0.0001.

### Determination of ATP content

NAC pre-incubation rescued the cells from the detrimental effects of silver disruption of the TCA cycle. Next, the downstream effect of TCA cycle salvage by NAC was measured in terms of ATP production to demonstrate the rescue of respiration in these cells (Figure 8). Cells pre-incubated with 10 mM NAC demonstrated significantly higher ATP production upon exposure to silver compared with cells exposed to silver alone (*p* ≤ 0.0001). Thus, NAC pre-incubation rescued cells from disruption of the TCA cycle and electron transport chain to maintain ATP production at a comparable rate to the control group.

**Figure 8.**
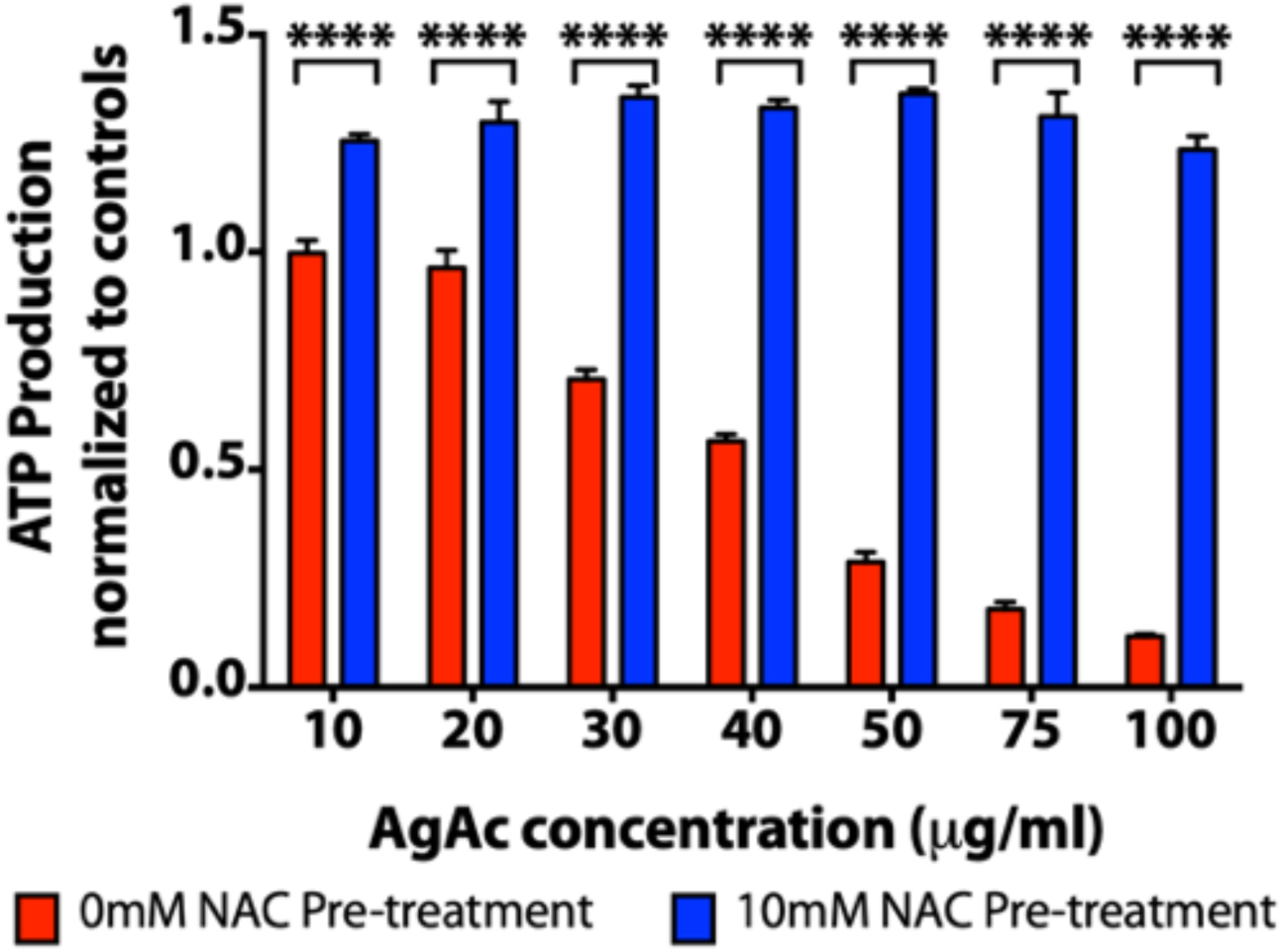
ATP production in human bronchial epithelial (16HBE) cells pre-incubated with 0 or 10 mM N-acetyl cysteine (NAC) followed by exposure to silver acetate for 8 h. ****: *p* ≤ 0.0001.

### Antimicrobial activity of silver with or without NAC pretreatment

Antimicrobial activity of silver acetate with or without pre-incubation with NAC was measured using a standard CLSI broth-microdilution method (Table 1). The minimum inhibitory concentration (MIC) of silver acetate did not change when the bacteria were pre-incubated with 0 or 10 mM NAC, demonstrating the selectivity of NAC to rescue eukaryotic cells without altering its antimicrobial activity.

**Table 1.**
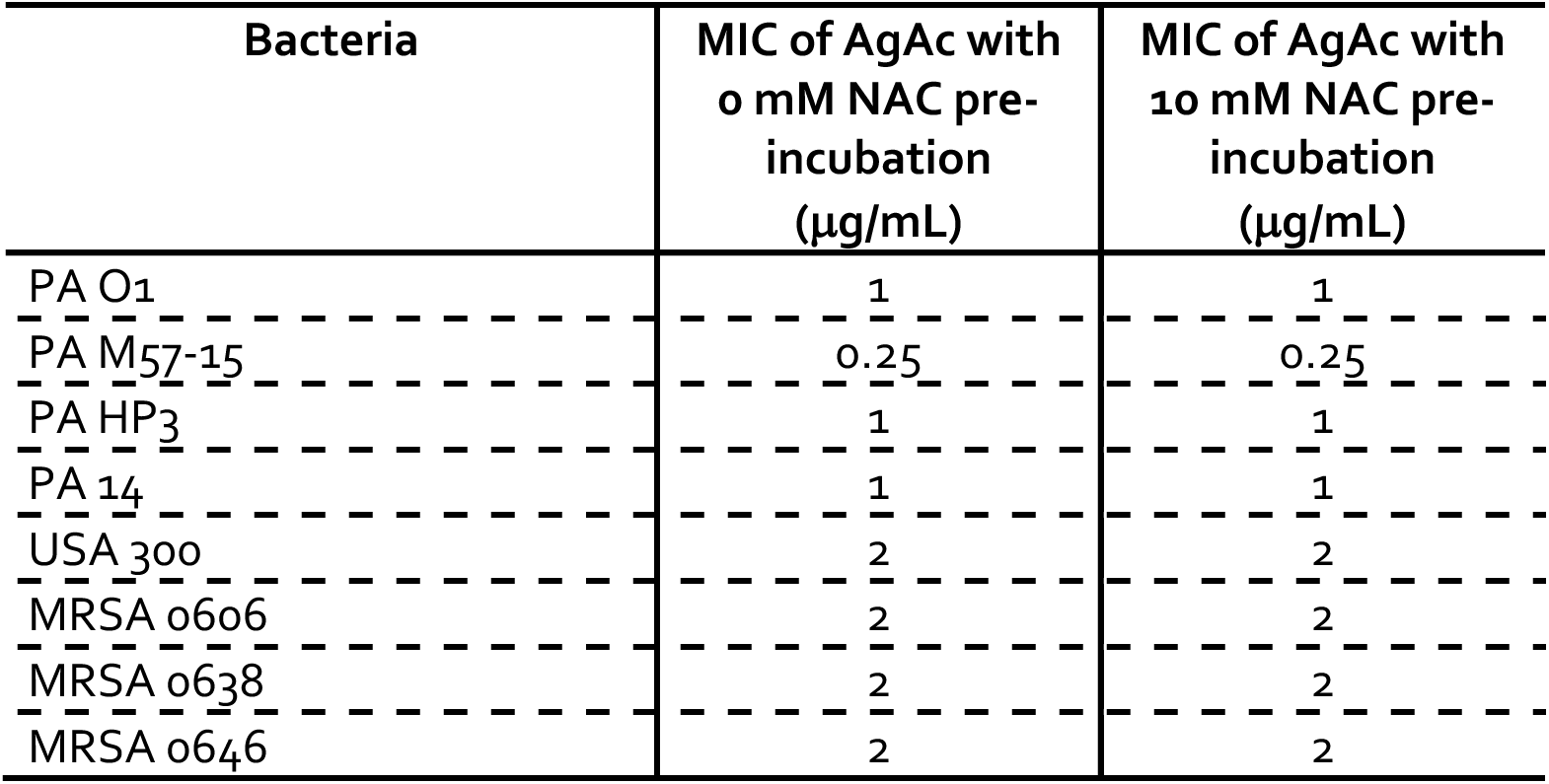
Minimum inhibitory concentration (MIC) of silver acetate (AgAc) against laboratory and clinical isolates of *P. aeruginosa* and MRSA upon 2 h pre-incubation with 0 or 10 mM NAC.

## Discussion

The multiple mechanisms of action that make silver an attractive broad-spectrum antimicrobial may also contribute to its toxicity towards eukaryotic cells. Several reports have demonstrated that exposure to silver at low concentrations results in apoptosis, and exposure at high concentrations results in necrosis (32, 35). Cell death, via apoptosis or necrosis, after silver exposure is often a result of ROS generation (32).

Mitochondria are considered the primary sources of ROS (30, 36). Generation of ROS in the mitochondria is a tightly regulated process that relies on the electron transport chain (ETC) for ROS formation and cellular ROS scavengers such as enzymes, superoxide dismutase (SOD), glutathione peroxidase (GP), thioredoxin (TRX), and peroxiredoxin (PRX), as well as antioxidants such as, glutathione, for countering the generated ROS (30). Under physiological conditions, glucose is converted to pyruvate during glycolysis, yielding two molecules of ATP, the energy currency in the cell (37). The pyruvate molecules are then transported into the mitochondria, where they are further broken down by the citric acid cycle to yield 36 molecules of ATP and ROS (37). Until recently, ROS was generally regarded as a byproduct of this process. However, in the last two decades, normal ROS levels have been implicated as second messengers in signal transduction pathways (38). For example, mitochondrial ROS (mtROS) plays a critical role in regulation of apoptosis, stem cell differentiation, autophagy, as well as cellular and tissue level inflammation (39). Upon exposure to chemicals including Ag^+^ ions, an imbalance of ROS generation results in oxidative stress causing damage to all three classes of biologic macromolecules - lipids, proteins, and nucleic acids (36, 38). High levels of oxidative stress induce cell death via one of two pathways: apoptosis or programmed necrosis (40).

Our results demonstrate a significant increase in the ROS and superoxide levels in human cells after exposure to silver (Figure 4). The downstream effects of ROS are complex, including activation of several pathways that ultimately lead to cell death (30, 36, 40). Mitochondrial DNA is highly susceptible to oxidative damage by mtROS due to its close proximity and a lack of protection by histones (36). To that end, the level of oxidatively modified bases in mtDNA has been found to be 10-20 fold higher than in nuclear DNA (36). In addition, nucleotide binding domain-leucine rich repeat (NLR) proteins recognize damage associated molecular patterns (DAMPs) in the mitochondrial DNA to initiate pyroptosis, a mode of induced cell death (39). Concurrently, mtROS stimulates lipid peroxidation, ultimately leading to disruption of mitochondrial membrane potential and mitochondrial calcium buffering capacity (36). ROS mediated opening of Ca^2+^ dependent mitochondrial permeability transition (MPT) pore results in ATP depletion, ultimately leading to necrosis (36, 41). MPT induction in part of the mitochondrial population, leaving the unperturbed mitochondria to produce sufficient ATP, results in caspase mediated apoptosis (36). These observations are consistent with our results; exposure to silver results, ROS induced oxidative stress leading to depletion of ATP (Figure 8) and cell death.

The surplus of superoxide and ROS interacts with critical proteins also resulting in activation of pathways that lead to cell death. Superoxide damages and deactivates iron-sulfur proteins such as aconitases (30). Superoxide rapidly reacts with the [4Fe-4S] to yield a catalytically inactive [3Fe-4S] protein, ultimately blocking the conversion of citrate to isocitrate by aconitases in the TCA cycle (30, 36). These effects are further exacerbated by the production of Fe^2+^ and hydrogen peroxide upon interaction between superoxide and thiol containing proteins. Released hydrogen peroxide can cause further oxidative damage to DNA, lipids, and proteins (36, 38). This inhibition of aconitases after exposure to ROS is typically associated with accumulation of citrate (30); however, our results show a reduction in citrate levels. This depletion of citrate can be attributed to the inactivation of pyruvate dehydrogenase kinase after exposure to mtROS (30). Pyruvate dehydrogenase kinase 2 (PDHK2), a major player of the pyruvate dehydrogenase complex (PDC) is responsible for catalytic conversion of pyruvate into acetyl-coA, a precursor of citrate in the TCA cycle. The mechanism of deactivation of these enzymes is not limited to the mitochondrial ROS; these proteins, including pyruvate dehydrogenase (PDH), can also be inhibited upon interaction with metal ions (42). Samikkannu *et al.* (42) have demonstrated inhibition of PDH after interaction with metal ions, as well as ROS. Thus, the combined deactivation of these two enzymes results in depletion of acetyl-coA, citrate, and isocitrate levels, key metabolites for the TCA cycle, ultimately resulting in disruption of the TCA cycle. The downstream effects of the loss of citrate directly impacts the TCA cycle, as demonstrated by reduced levels of downstream metabolites, glutamate, aspartate, fumarate, and malate (Figure 7). In response to this breakdown, there is a significant reduction in the ATP produced during the process, after exposure to silver (Figure 8).

The effect of silver on mitochondria is one of the primary mechanisms of silver toxicity. This initial insult to the mitochondria initiates several downstream signaling pathways that contribute to silver toxicity. For instance, DNA damage caused by ROS insult induces p53 stabilization (36, 43). Subsequent translocalization of p53 to the mitochondria triggers expression of the Bcl-2 family proteins Bax and PUMA, release of cytochrome c and activation of the caspase cascade (36). Similarly, thiol residues on several cysteine containing proteins act as a switch, which upon interaction with ROS can result in activation or repression of multiple transcription factors. These thiol based switches act by either inhibiting enzymatic activity or inducing conformational changes to regulate transcription factors (30). Further, TP53-induced glycolysis and apoptosis regulator (TIGAR), upregulated by p53, inhibits glycolysis, and shifts the glucose flux into the pentose phosphate pathway (PPP) (36). Similarly, inhibition of pyruvate kinase isoform (PMK2) forces activation of the PPP (30). This shift to the PPP, which is responsible for generating majority of the NADPH, is key defense mechanism activated by the cell in response to the oxidative damage. The products of the PPP are subsequently shuttled back into the glycolytic pathway. In addition to activation of the PPP, PMK2 is also thought to play a critical role in the antioxidant defense system by diverting 3-phosphoglycerate to glutathione synthesis, via the phosphoserine synthesis pathway (30, 44). Activation of this pathway should result in an increase in the reduced glutathione (GSH) levels, which further deactivates ROS to form the oxidized glutathione (GSSG) (38). Our results indicate a reduction in the reduced form of glutathione after exposure of human cells to silver (Figure 5) and are consistent with the results obtained by Arora *et al.* after exposing cells to silver nanoparticles (32). However, the expected increase in the GSSG levels is not observed. This unexpected behavior is likely caused by the high affinity of silver ions towards the thiol groups present on glutathione. The silver glutathione complex blocks the ability of GSH to neutralize ROS. Thus, silver associated toxicity is a direct consequence of a combination of silver interaction with cellular components and ROS generation. These two processes are closely related, often complementary, and their individual effects cannot be separated. The downstream response of all these mechanisms results in a Warburg-like effect, where the glucose consumed by the cells is not completely oxidized and is secreted as lactate (Figure 6). For use of silver as an effective antimicrobial, we must address all of these toxicity mechanisms without altering its antimicrobial activity.

The multiple cell-death mechanisms activated by ROS generated by exposure to silver have garnered attention of several researchers (28, 29, 32, 34). Antioxidants have been explored to check the ROS generated after exposure to metal (31). N-acetyl cysteine (NAC) is one of the most common antioxidants investigated to abrogate the toxicity of silver, as well as other metal ions (29, 33–35, 45). Several groups have demonstrated successful rescue of mammalian cells upon pre-exposure to NAC followed by exposure to metal ions (33, 35, 44, 45). NAC pre-incubation results in significant ROS reduction upon exposure to silver (Figure 4). However, it is unclear if the reduction in the ROS levels is a direct consequence of the ROS scavenging activity of NAC or a downstream effect of silver-NAC interactions. In order to elucidate the mechanism of rescue and develop a future potential therapeutic that can ameliorate toxicity of silver-based therapeutics such as, silver antimicrobial bandages, we investigated several antioxidants and their ability to rescue mammalian cells from silver toxicity. Of all the evaluated antioxidants, NAC is the only antioxidant that demonstrates significant rescue (Figure 3, Supplementary Fig. S1, S2, and S3). Melatonin, which demonstrates only slightly lower antioxidant capacity compared with NAC, does not result in rescue of cells from silver associated toxicity. Although ascorbic acid has a higher antioxidant capacity than NAC, ascorbic acid failed to rescue the cells from silver toxicity. Indeed, among the antioxidants tested, NAC exclusively rescues cells from silver associated toxicity; the ROS levels in cells pre-incubated with NAC followed by up to 50 μg/mL silver are similar to the negative controls, cells that received neither NAC nor silver. The failure of ascorbic acid to rescue silver-exposed cells suggests that antioxidant activity is not the only mechanism through which NAC pre-treatment resulted in lower ROS levels.

Unique structural properties position NAC to abrogate the multiple toxicity mechanisms induced by silver cations. NAC is a precursor for glutathione, however, the glutathione levels of cells incubated with NAC are similar to those of control cells. Thus, the reduction in ROS is not directly influenced by glutathione. We hypothesized that the thiol groups present on NAC bind with silver, which would prevent it from binding with DNA, and generating ROS. We confirmed the ability of NAC to bind with silver using ^1^H nuclear magnetic resonance (NMR) spectrometry (Figure S5). The hydrogen atoms associated with the primary carbon on NAC demonstrate a distinct shift (at 2-3 ppm) in the signal in the presence of silver. Further, NAC and silver self-assemble into a hydrogel, confirming the interaction between NAC and silver. Similar NAC-based hydrogels after interaction with silver, gold, and copper have been reported by Casuso *et al* (46). In addition, NAC is known as an antiapoptotic agent, and inhibits several apoptotic pathways including, NF-κB, MEK/ERK, and the JNK pathway, promoting cell survival (47, 48). The downstream effect of NAC treatment is underscored by the significant improvement in the metabolic state of the cell. Cells treated with NAC demonstrate normal levels of TCA cycle metabolites after silver exposure. These cells also demonstrate normal lactate production and ATP levels. The ability of NAC to rescue eukaryotic cells is not limited to bronchial epithelial cells; human dermal fibroblasts also demonstrate similar cell viability patterns. Finally, the ability of NAC to rescue eukaryotic cells from silver associated cell death does not affect the antimicrobial activity of silver; when *P. aeruginosa* and MRSA isolates were pre-exposed to 10 mM NAC, the MIC of AgAc was identical to that of NAC-free AgAc (Table 1). Moreover, NAC is frequently used as a mucolytic in patients suffering from cystic fibrosis (CF) (49), chronic obstructive pulmonary disorder (COPD) (50), and chronic bronchitis (51, 52). The benefits of NAC have led to its inclusion in The World Health Organization’s (WHO) list of essential medicines (53). Thus, a silver/NAC combination presents a unique therapeutic strategy that can effectively eradicate bacterial infections without causing toxicity to eukaryotic cells.

In conclusion, silver is a potent, FDA approved, antimicrobial that is widely used to eradicate infections associated with MDR pathogens. We report here a novel mechanism that results in rescue of mammalian cells from silver associated toxicity without altering its antimicrobial activity. Cells incubated with silver demonstrate high levels of ROS, which causes disruption of the TCA cycle and reduction in ATP production, ultimately leading to cell death via apoptosis or necrosis. On the other hand, cells pre-incubated with NAC followed by silver do not demonstrate signs of oxidative stress, show a normal metabolic state, as well as ATP production, which translates to lower silver toxicity. Thus, the silver/NAC combination has tremendous potential as a future therapeutic with potent antimicrobial activity with a large therapeutic window.

## Materials and Methods

### Reagents

Silver acetate, Dulbecco’s Modified Eagle’s Medium (DMEM) powder (without glucose, phenol red, L-glutamine, sodium pyruvate, and sodium bicarbonate), D-glucose, L-glutamine, sodium bicarbonate (NaHCO_3_), HEPES buffer, penicillin-streptomycin (100X stock), trypsin-EDTA solution, sodium hydroxide (NaOH, 1N), methanol, Minimum Essential Medium (MEM) with Earle’s Balanced Salts and non-essential amino acids, fetal bovine serum (FBS), and N-acetyl cysteine (NAC) were obtained from Sigma-Aldrich Corporation (St. Louis, MO). Uniformly labeled [U^13^C] glucose was obtained from Cambridge Isotope Laboratories, Inc. (Andover, MA). Opti-MEM (without phenol red), alamarBlue^®^ Cell Viability Kit (Cat # DAL1100), ATP Determination Kit (Cat # A22066), and Phosphate Buffered Saline (PBS) solution (10X) were obtained from Thermo Fisher Scientific, Inc. (Waltham, MA). Total Antioxidant Capacity Assay Kit (Cat # ab65329), Cellular ROS/Superoxide Detection Assay Kit (Cat # ab139476), GSH/GSSG Ratio Detection Assay Kit II (Cat # ab205811), Deproteinizing Sample Kit (Cat # ab204708), and Mammalian Cell Lysis Buffer 5X (Cat # ab179835) were purchased from Abcam (Cambridge, MA). Tissue culture flasks, tissue culture dishes (Φ = 60 mm), 24-well plates, 96-well plates, Tryptic soy agar (TSA) plates, and Mueller-Hinton (MH) broth were obtained from Becton Dickinson and Company (Franklin Lakes, NJ), respectively. Distilled deionized water (DH_2_O) was obtained from a Milli-Q biocel system (Millipore, Billerica, MA) and sterilized in an autoclave. All the above chemicals were used without further purification.

### Cell culture

Human bronchial epithelial cell line (16HBE140-) generously provided by Dr. D. Gruenert (University of California, San Francisco, CA) are a human bronchial epithelial cell line transformed with SV40 large T-antigen using the replication-defective pSVori plasmid (54). 16HBEs were used between passages of 20 and 40 for all experiments. 16HBE cells were cultured in Minimum Essential Medium (MEM) with Earle’s Balanced Salts and non-essential amino acids supplemented with 10% fetal bovine serum (FBS), 1% L-glutamine, and 1% penicillin-streptomycin (P/S) solution at 37°C in an incubator (5% CO_2_, 100% RH), unless otherwise noted. When the cells reached 90-95% confluency, they were harvested by trypsinizing and sub-cultured.

### Silver induction of reactive oxygen species (ROS) and superoxide

Cellular ROS and superoxide levels were measured in 16HBE cells using a Cellular ROS/Superoxide Detection Assay Kit according to manufacturer’s recommended protocol. Briefly, cells were seeded at a density of 25,000 cells/well in a black wall/clear bottom 96-well plate and incubated for 24 h as described above. Next, the feeding media was aspirated and cell were incubated with fresh media supplemented with or without 10 mM NAC for 2 h. Finally, the NAC solution was removed and cells were incubated with 100 μL of 0, 10, 20, 50, or 100 μg/mL silver acetate containing 1X ROS/Superoxide detection mix. Upon staining, the fluorescence signal from the two fluorescent dyes, green signal from ROS detection probe (Ex/Em = 490/525 nm) and orange signal from superoxide detection probe (Ex/Em = 550/620 nm), were quantified using a BioTek Instruments Cytation 5 Multimode Reader at 0, 4, 6, 8, and 24 h. The fluorescence signal was normalized to the drug free controls (0 mM NAC + 0 μg/mL silver acetate). All experiments were performed with 6 technical replicates and a minimum of 2 biological replicates.

### Activity of antioxidants

Antioxidant activity of NAC, ascorbic acid, and melatonin was measured using a Total Antioxidant Capacity Assay Kit according to manufacturer’s recommended protocol. A standard curve correlating the Trolox concentration to the antioxidant capacity was generated according to manufacturer’s protocol. NAC, ascorbic acid, and melatonin were dissolved at 10mM concentration in distilled – deionized water (d. d. water) and serially diluted. All experimental and standard solutions were protected from light, incubated with colorimetric Cu^+2^ probe for 1.5h with constant shaking and absorbance was measured at 570 nm using a BioTek Cytation 5 Multimode Reader. The antioxidant capacity of test solutions, NAC, ascorbic acid, and melatonin were then correlated to the standard curve and presented as a function of the final Trolox concentration. Next, the antioxidant capacity of NAC, ascorbic acid, and melatonin pre-incubated 16HBE cells was also measured. Two million 16HBE cells were seeded into each well of a 12-well plate and incubated overnight as described above. The feeding media was then replaced with fresh feeding media or media containing 10 mM NAC, ascorbic acid, or melatonin. After a 2 h incubation with the antioxidants, cells were washed with cold PBS, re-suspended in 100 μL d. d. water, homogenized by pipetting, and incubated on ice for 10 min. Finally, the insoluble cell debris was removed by centrifugation and the supernatant analyzed as described above to determine the total antioxidant capacity. All experiments were performed with 4 technical replicates and two biological replicates.

### Abrogation of silver acetate toxicity through pre-incubation with antioxidants

Toxicity of silver acetate with or without pre-incubation with antioxidants was assessed on 16HBE cells using an alamarblue® Cell Viability Assay according to manufacturer’s recommended protocol. Cells were seeded at a density of 25,000 cells/well in a 96-well plate and incubated overnight as described above. At 24 h, media was aspirated, and cells were pre-incubated with 80 µL of 0, 0.01, 0.1, 1.0, 2.5, 5, 7.5, and 10 mM concentrations of NAC, ascorbic acid, and melatonin for 2 h. Next, the antioxidant supplemented media was replaced with 100 µL feeding media containing 0, 10, 20, 30, 40, 50, 75, and 100 µg/mL silver acetate. Alamarblue test reagent was added to each well, and the plates were incubated as described above. At 8 and 24 h timepoints, absorbance was measured at 570 and 600 nm, normalized to media only controls, and analyzed per manufacturer’s instructions. All experiments were performed with 6 technical replicates and 3 biological replicates. These results were verified using an CyQUANT® Cell Proliferation Assay Kit (Supplementary Information).

### Glutathione concentrations after pretreatment with NAC

Glutathione levels in 16HBE cells pre-incubated with NAC were determined using a GSH/GSSG Ratio Detection Assay Kit II according to manufacturer’s recommended protocol. Five million 16HBE cells were seeded in each well of a 6-well plate as described above. At 24 h, the feeding media was replaced with fresh feeding media supplemented with or without 10 mM NAC and incubated for 2 h. Next, cells were incubated with 0, 10, and 100 µg/mL silver acetate for 1 h and glutathione levels measured. Briefly, cells were washed with cold PBS, re-suspended in 300 µL ice cold Mammalian Cell Lysis Buffer and homogenized by pipetting. The cell lysate was then centrifuged to remove the cell debris and the supernatant was carefully collected and deproteinized using a Deproteinizing Sample Kit. The deproteinized samples were then diluted using lysis buffer, mixed with glutathione (GSH) and total glutathione (TGAM or GSH + GSSG) assay probes, incubated for 60 minutes protected from light, and fluorescence signal measured at Ex/Em = 490/520 nm using a BioTek Cytation 5 Multimode Reader. The fluorescence signal from the experimental values were then correlated to the glutathione (GSH and GSH + GSSG) standard curves generated to determine the intracellular glutathione concentrations. Experiments were performed with 4 technical replicates and 3 biological replicates.

### Analysis of total metabolite pool size and metabolite labeling patterns using Gas Chromatography-Mass Spectroscopy

16HBE cells were seeded at a density of 250,000 cells per dish in a 60mm cell culture dish and incubated until they reached 90% confluency. During this period, feeding media was replaced every 48h. Once confluent, the media was aspirated, cells were washed with 1X PBS, and incubated with 2 mL 0 or 10 mM NAC for 2 h. Next, the NAC supplemented media was replaced with 2 mL 4mM GLN-10mM D-[U^13^C]-GLC medium containing 0, 10, 20, 30, 40, 50, 75, and 100 µg/mL silver acetate. At 8 h, the feeding medium from each plate was collected, centrifuged at 1000 rpm for 5 min to remove any cell debris, and frozen at −80°C, until further analysis. The cells were washed twice with 1X PBS, re-suspended by gentle scraping in 1 mL chilled 50% methanol solution, cell lysate collected in centrifuge tubes, flash frozen using liquid nitrogen, and stored at −80°C till analysis.

The supernatant obtained from cells pre-treated with or without NAC and exposed to various concentrations of silver acetate in 4 mM GLN-10 mM D-[U^13^C]-GLC medium (all time points) was thawed and analyzed for concentrations of glucose and lactate using a BioProfile BASIC Analyzer (Nova Biomedical, Waltham, MA). MEM and stock solution of 4 mM GLN-10 mM D-[U^13^C]-GLC medium treated in an identical manner were used as controls.

Cell suspensions frozen in 50% methanol were thawed and subjected to three additional freeze-thaw cycles using liquid nitrogen and a water bath. Subsequently, the cell suspensions were centrifuged at 14,000 rpm for 10 min to remove cell debris, and the supernatants were transferred to individually labeled glass drying tubes. 10 μl of an internal standard (50 nmols of sodium 2-oxobutyrate) was added to each tube at this time, and the samples were air-dried on a heat block. The dried samples were derivatized by addition of 100 μl of Tri-sil HTP reagent (Thermo Scientific) to each tube, capping the tube, vortexing the samples, and placing them on the heat block for an additional 30 min. The derivatized samples were transferred to auto-injector vials and analyzed using gas chromatography-mass spectroscopy (GC-MS; Agilent Technologies, Santa Clara, CA). Separately, the cell pellets with residual cell lysate was collected, contents thoroughly mixed with 200 μL 0.1 N sodium hydroxide, and heated to 100°C to extract and solubilize the proteins. The samples were cooled and analyzed using a standard BCA assay to quantify the protein content. All metabolite concentrations determined using the BioProfile Basic-4 analyzer (NOVA) and GC-MS were normalized with the protein content.

### Determination of ATP content

ATP production by 16HBE cells with and without pre-incubation with NAC followed by incubation with silver acetate was determined using an ATP determination kit using manufacturer’s recommended protocol. 50,000 16HBE cells were seeded in each well of a 96-well plate and the cells were allowed to attach. At 24 h, media was aspirated, and cells were pre-incubated with 80 µL of 0 or 10 mM concentrations of NAC for 2 h. Next, the antioxidant supplemented media was replaced with 100 µL feeding media containing 0, 10, 20, 30, 40, 50, 75, and 100 µg/mL silver acetate. Following an 8 h incubation with silver acetate, the media was aspirated, cells were washed with 100 µL 1X PBS, and incubated with 100 µL lysis buffer for 15 min. Finally, the cell lysate was collected and the ATP concentration determined and correlated to an established standard curve. Briefly, a standard reaction mixture consisting of molecular grade water, reaction buffer, Dithiothreitol (DTT) solution, D-luciferin, and firefly luciferase at manufacturer recommended concentrations was prepared. Next, 10 µL of standard ATP solution or cell lysate was mixed with 90 µL standard reaction mixture in a 96-well white bottom plate and luminescence was measured at 560 nm. Background luminescence was subtracted from all readings and the data were normalized to drug free controls.

### Antimicrobial activity of silver

Antimicrobial activity of silver was evaluated against laboratory and clinical isolates of *Pseudomonas aeruginosa* (PA O1, PA M57-15, PA HP3, and PA14) as well as methicillin-resistant *Staphylococcus aureus* (MRSA; USA 300, MRSA 0606, MRSA 0638, and MRSA 0646), with or without pre-incubation with NAC. Frozen stocks of bacteria were struck onto TSA plates and allowed to grow for 18-24 h at 37°C. A single colony was used to inoculate 5 mL MH broth and grown to an OD_650_ = 0.40 at 37°C on an orbital shaker. Next, the bacteria were centrifuged at 2500 rpm for 15 m at 4°C, supernatant aspirated, and bacterial pellets were re-suspended in 2 mL MH broth supplemented with 0 or 10 mM NAC. Bacterial suspension was then incubated at 37°C with orbital shaking for 2 h, centrifuged again to remove the NAC solution, and re-suspended in NAC free MH broth to OD_650_ = 0.4. Finally, minimum inhibitory concentrations (MIC) against silver acetate were determined using standard Clinical and Laboratory Institute (CLSI) broth-microdilution method. Briefly, bacterial suspension at a concentration of 5E5 colony forming units (CFU) per milliliter was incubated with a silver acetate at a final concentration of 0.13, 0.25, 0.5, 1, 2, 4, 8, 16, and 32 μg/mL silver acetate at 37°C for 18-24h, under static conditions. The MIC was determined as the lowest concentration resulting in no bacterial growth upon visual inspection. All experiments were performed in triplicate.

### Statistics

All data were analyzed using GraphPad Prism 7 (GraphPad Software, Inc., La Jolla, CA). A two-way analysis of variance (ANOVA) followed by a *post hoc* Sidak’s or Tukey’s test with multiple comparisons between means at each concentration of silver acetate was used to determine the significant difference. Additionally, non-linear regression was used to deduce the lethal dose at median cell viability (LD_50_) for cell viability assays. A *p* ≤ 0.05 was considered significantly different.

---

**Figure S1.**
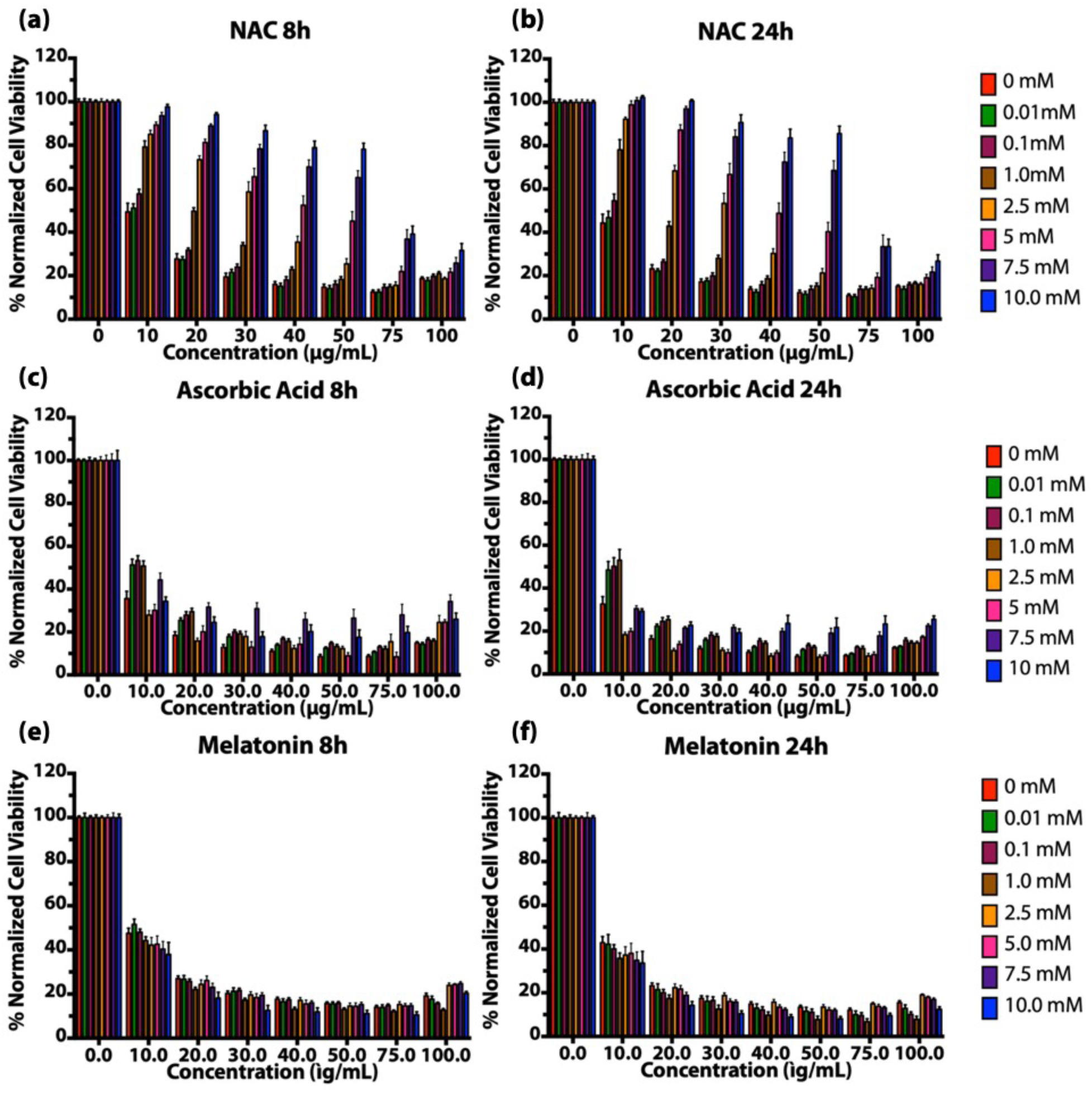
Viability of human bronchial epithelial (16HBE) cells upon pre-incubation with (a) NAC, (c) ascorbic acid, and (e) elatonin for 2 h followed by silver acetate exposure for 8 h, or (b) NAC, (d) ascorbic acid, and (f) melatonin for 2h followed by silver acetate exposure for 24 h.

**Figure S2.**
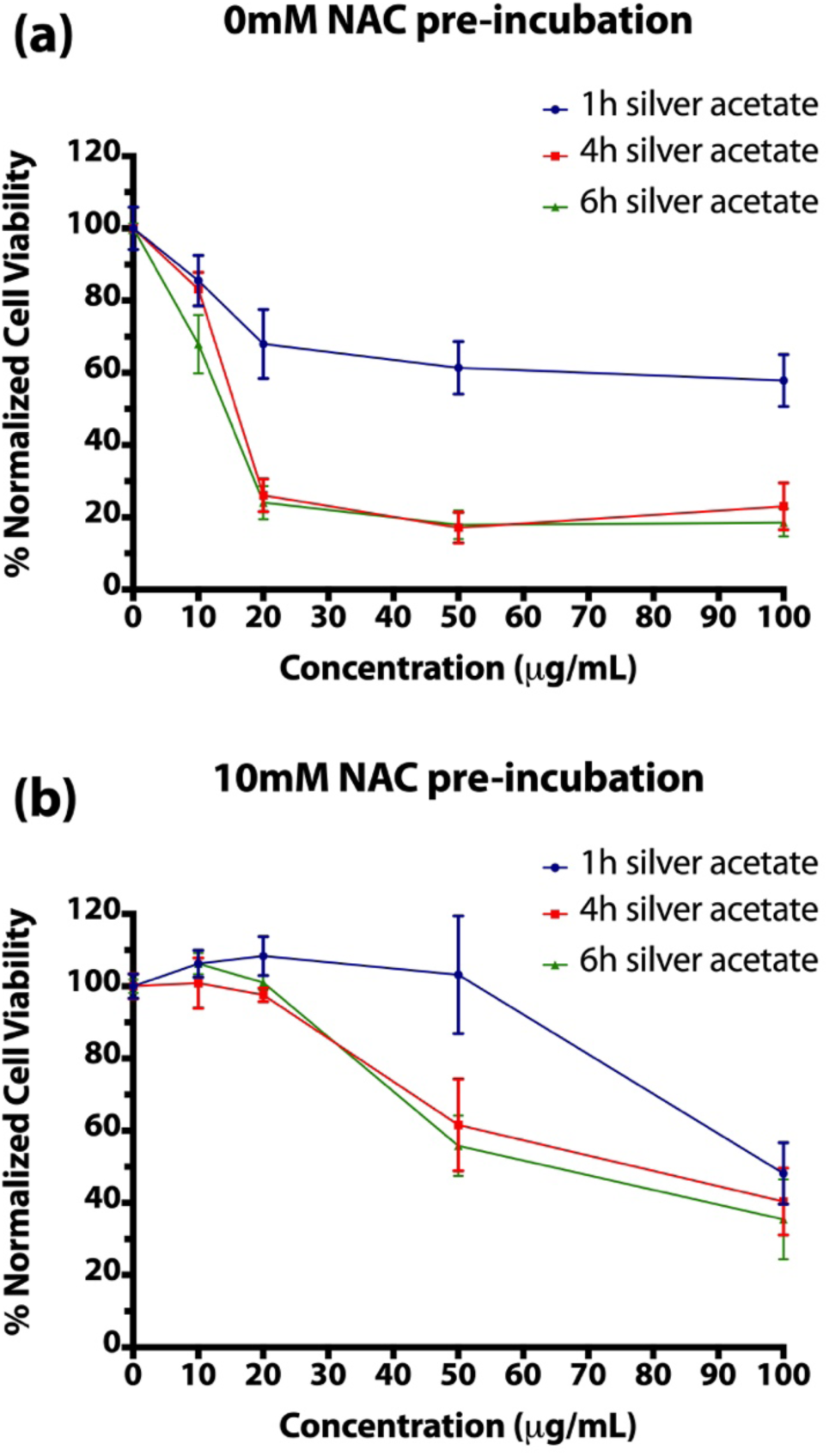
Viability of human bronchial epithelial (16HBE) cells upon pre-incubation with (a) 0 or (b) 10 mM NAC followed by silver acetate exposure for 1, 4, and 6h measured using a CyQUANT® Cell Proliferation Assay Kit.

**Figure S3.**
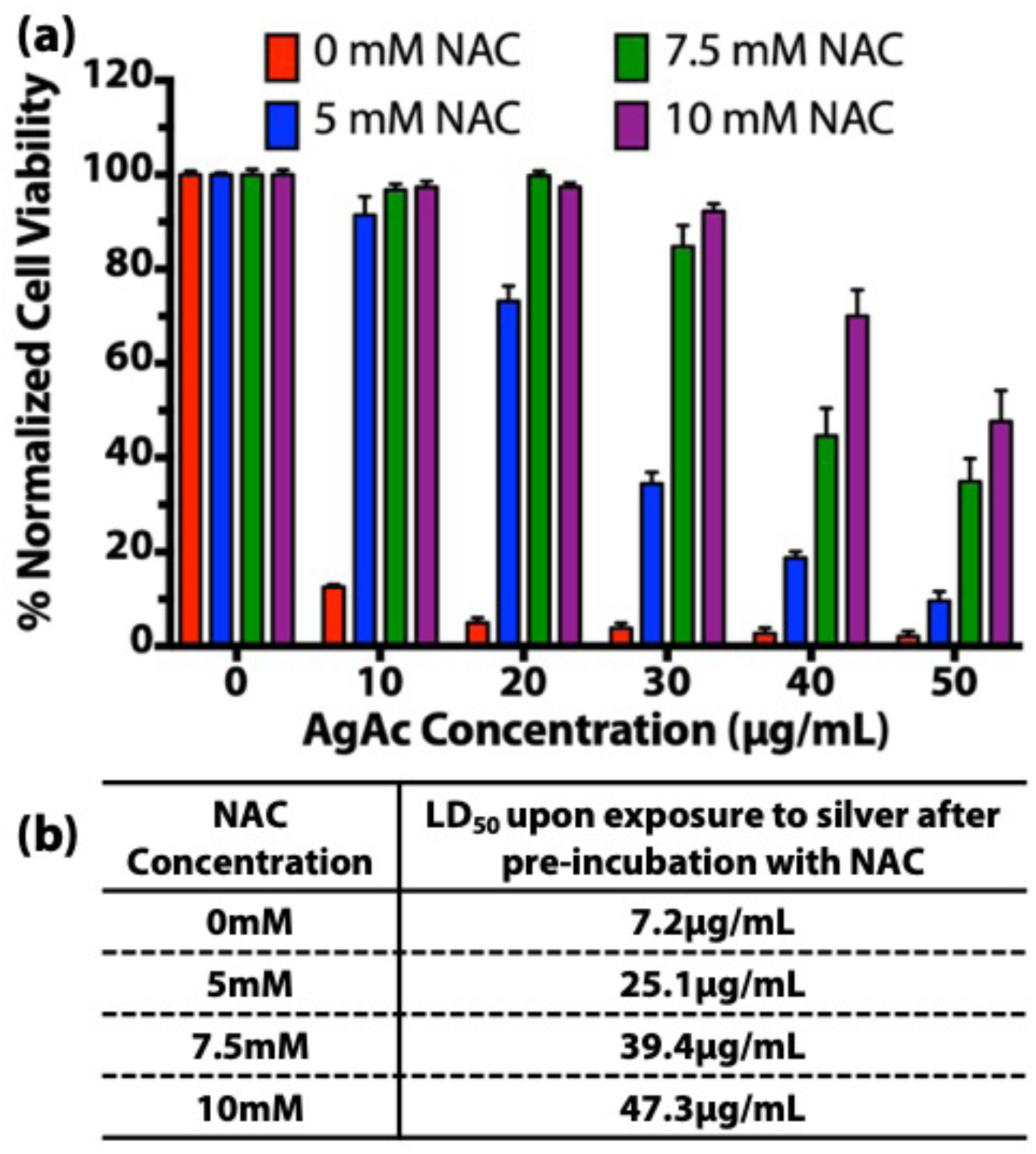
(a) Viability and (b) lethal dose at median cell viability (LD_50_) of human dermal fibroblasts (HDF) cells upon pre-incubation with 0, 5, 7.5, or 10 mM NAC followed by exposure to silver acetate for 24 h.

**Figure S4.**
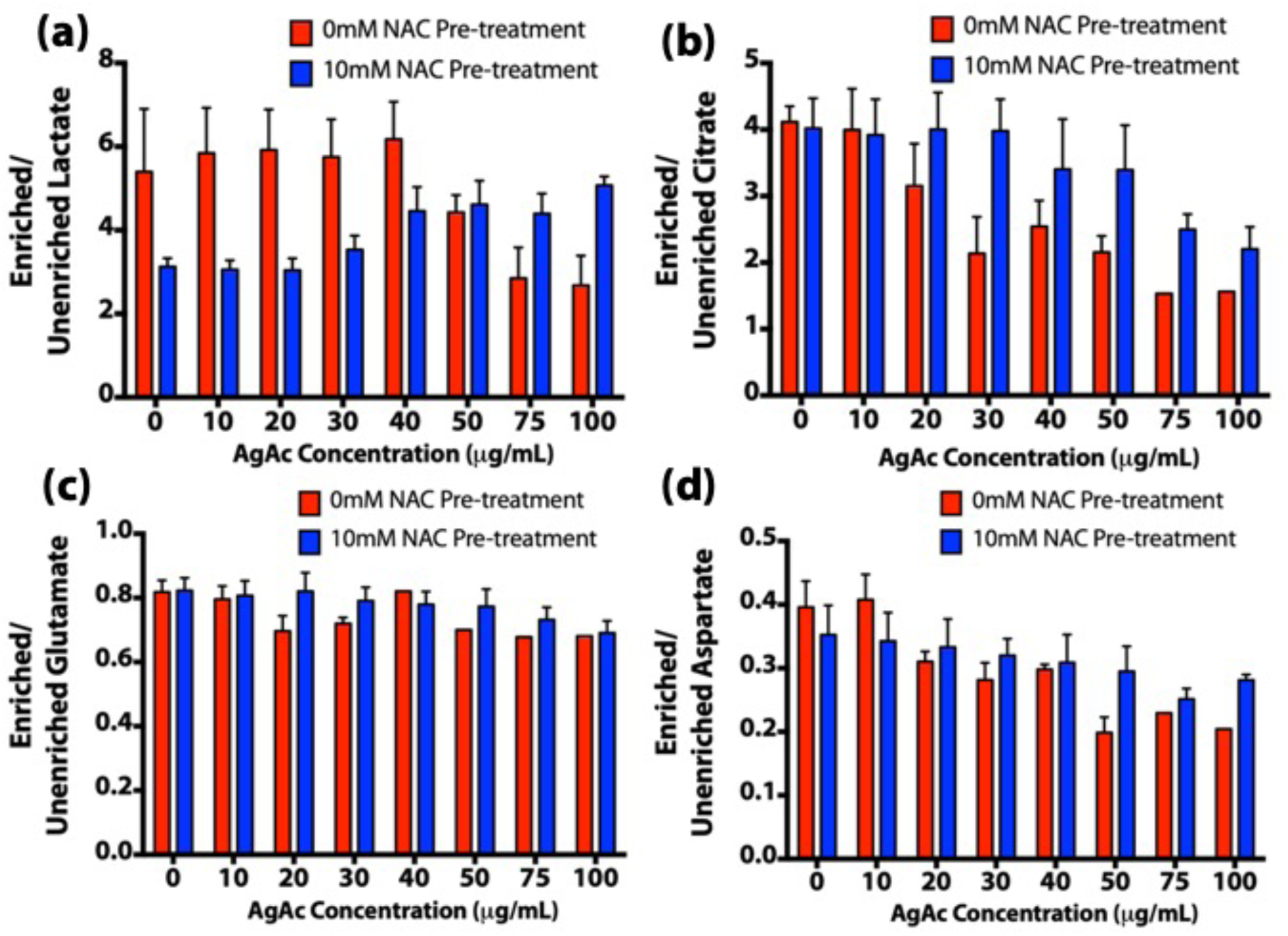
Labeling patterns for key citric acid cycle metabolites, (a) lactate, (b) citrate, (c) glutamate, and (d) aspartate with or without silver acetate.

**Figure S5.**
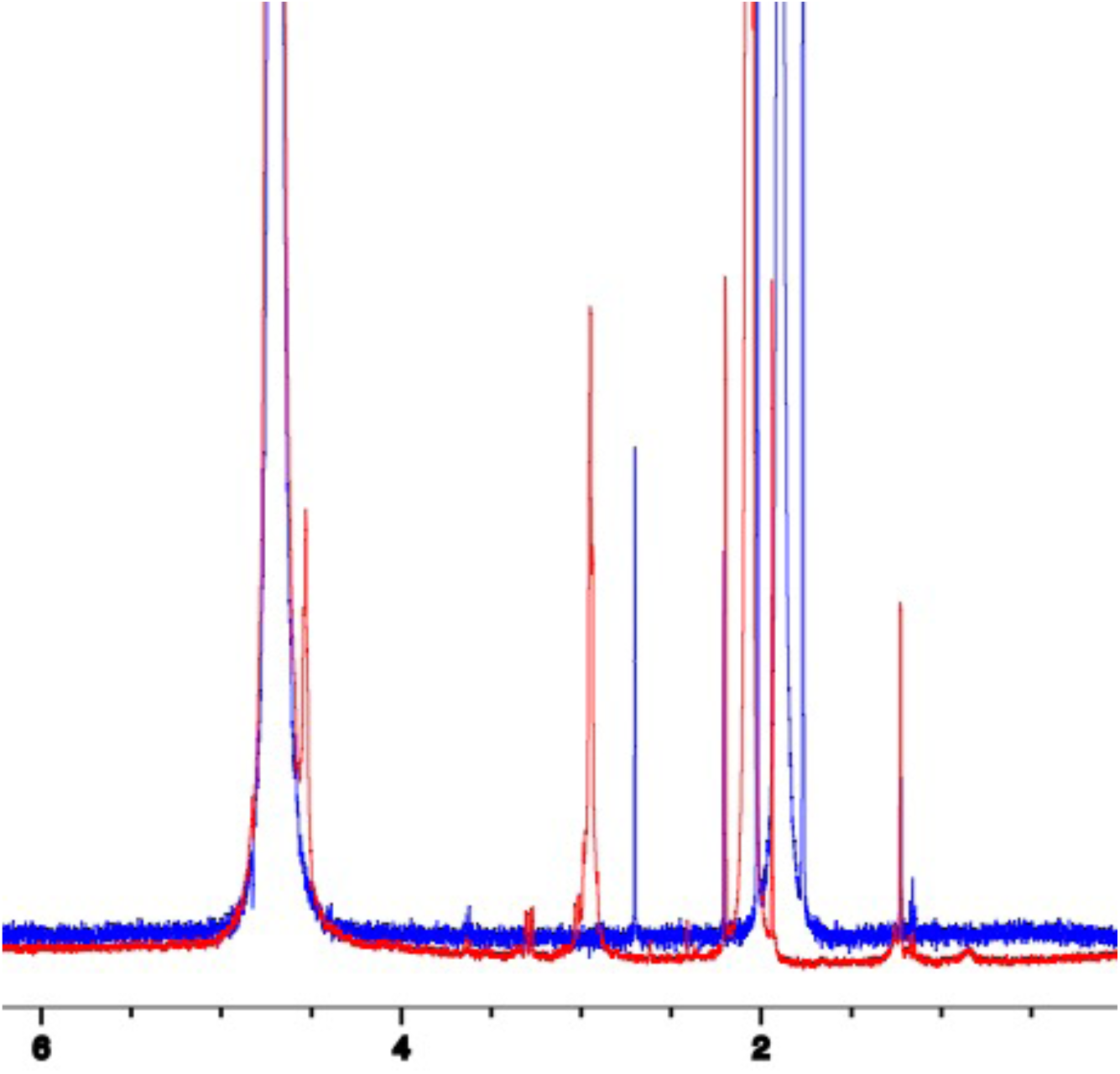
^1^H Nuclear magnetic resonance (NMR) of N-acetyl cysteine (blue) and a combination of silver acetate and N-acetyl cysteine (red) dissolved in deuterated water.

